# Molecular evolution and genetic analysis of Mesta yellow vein mosaic virus and associated betasatellites

**DOI:** 10.1101/2021.09.05.459025

**Authors:** Rasel Ahmed, Rajnee Hasan, Md. Wali Ullah, Borhan Ahmed

**Author notes:** **Correspondence:** Rasel Ahmed, Basic and Applied Research on Jute Project, Bangladesh Jute Research Institute, Manik Mia Avenue, Dhaka-1207, Bangladesh.

## Abstract

Mesta yellow vein mosaic disease (MYVMD), one of the major diseases circulating mesta growing regions of Indian sub-continent, is responsible for serious yield loss in mesta crops. A complex of monopartite begomovirus, Mesta yellow vein mosaic virus (MYVMV) and associated betasatellite, is reported in several studies as the causal agent of MYVMD. However, all-inclusive molecular evolutionary analysis of so far available MYVMVs and associated betasatellites disseminating in this region is still lacking. In this study, by estimating and analyzing various indexes of population genetics and evolutionary parameters, we discussed the sources of genetic variations, population dynamics and different forces acting on the evolution of MYVMVs and associated betasatellites. The study finds recombination as a vital force in the evolution and diversification of begomovirus complexes in different geographic locations however, betasatellites were found to be exposed to more diverse recombination events compared to MYVMVs. Indian isolates are reported to have high frequency of polymorphism in this study which suggests a balancing selection or expansion occurring in Indian populations of begomoviruses. Higher degree of genetic differentiation and lower rate of gene flow calculated between the viral populations of Bangladesh and Pakistan is justified by the relatively far geographical distance between these two countries. Although the study detects overall purifying selection, the degrees of constraints acting on individual gene tested are found different. Coat protein (AV1) is estimated with very high nucleotide substitution rate which is very likely to result from the strongest purifying selection pressure (*d_N_/d_S_* = 0.131) calculated in this study on coat protein. The findings of this study on different evolutionary forces that shape the emergence and diversification of MYVMVs and associated betasatellites may provide directions towards future evolutionary trend analysis and development of comprehensive disease control strategies for begomoviruses.

## Introduction

Mesta is one of the cheapest and important bast fiber producing crops cultivated largely in Indian sub-continent. For being biodegradable and renewable in characteristics, the bast fibers are drawing remarkable attention worldwide to address several environmental issues. Two species of mesta namely *Hibiscus cannabinus* (kenaf) and *Hibiscus sabdariffa* (roselle) of the malvaceous family are cultivated mainly for producing lingo-cellulosic bast fibers which are used to manufacture traditional packaging materials such as rope, sacking, canvas, cordage, carpet backing etc. Beside this, the leaves and the seeds of both species are being used as vegetable and rich source of oil respectively and for having lower proportion of unsaturated fatty acids, the oil has potential medicinal value [1,2]. Moreover, mesta, particularly kenaf (*Hibiscus cannabinus*) is now gaining global interest as a non-woody source of quality pulp for tree-free newspaper industry.

The cultivation of mesta crops is highly susceptible to a disease called mesta yellow vein mosaic disease (MYVMD) which is alarmingly affecting and spreading the large mesta growing area in the sub-continent. The MYVMD is characterized by a typical ‘‘yellow vein network’’ showed in the leaf veins followed by chlorosis of the leaves. The bast fiber yield of the diseased plants is seriously affected due to the stunted growth, complete chlorosis and defoliation of the leaves in the advanced stage of the disease [3,4]. The disease was documented first time in West Bengal state of India which is located adjacent to Bangladesh [5]. Several districts, particularly Kishoreganj in Bangladesh also came under sporadic attack of yellow mosaic few years ago and the affected areas are increasing in a yearly basis (http://www.newagebd.net/article/18783/yellow-mosaic-virus-attacks-jute-crop). A monopartite begomovirus, Mesta yellow vein mosaic virus (hereafter called MYVMV for all the Mesta yellow vein mosaic viruses isolated from mesta plants) and an isolate of Cotton leaf curl Multan betasatellite are reported to be associated with MYVMD in eastern India [6,7]. A similar disease of mesta was also found to be associated with a monopartite begomovirus and a betasatellite in the northern part of India [3]. Some mesta leaf samples from northern India showing yellow mosaic symptoms were carried out for diagnosis and the identified virus particles showed similarity with another reported begomovirus Mesta yellow vein mosaic Bahraich virus and an isolate of Ludwigia leaf distortion betasatellite [8].

Begomovirus (transmitted by whiteflies, *Bemisia tabaci*) is the largest genus under the family Geminiviridae and are widely distributed geographically. Begomoviruses have either monopartite (DNA-A-like component) or bipartite (DNA-A and DNA-B) genomes [9] containing circular, single-stranded DNA (ssDNA) of approximately 2.7 Kbp. Typically, DNA-A has six open reading frames (ORFs) of AV1(coat protein), AV2 (pre-coat protein), AC1 (replication initiation protein or rep protein), AC2 (transcriptional activator protein), AC3 (replication enhancer protein) and AC4 (AC4 protein). Among these 6 ORFs AV1 and AV2 are on the virion-sense strand and rest are on the complementary-sense strand. Monopartite begomoviruses are often co-existed with satellite termed as betasatellites [9]. These betasatellite molecules depend on the helper begomoviruses for their replication and are approximately 1.3 Kbp in size containing one major ORF (βC1) on complementary strand. The βC1 ORF has been characterized to be involved in virus movement, suppressing host antiviral silencing as a pathogenicity determinant [10,11] and symptom induction [12]. In evolutionary aspects, nucleotide substitution and recombination rate in the ssDNA viruses are quite similar to that of RNA viruses [13]. Recombination is recognized as a vital process in the evolution of begomoviruses [14,15,16] and some studies in begomoviruses found high rate of recombination within recombination hotspots [17,18,19]. On the other hand, compound microsatellites (cSSRs) are reported to be potentially linked with the recombination of begomoviruses [20,21]. The simple sequence repeats or microsatellites (SSRs) are tandem repeats of short DNA sequences and considered as cSSRs when two or more closely located SSRs are separated by non-repeat sequences. The microsatellites have been extensively studied in viruses, prokaryotes, and eukaryotes [22,23,24] and found to be involved in gene regulation and transcription processes [25,26].

In this study, we collected kenaf leaf samples with yellow mosaic symptoms from different regions of Bangladesh and identified the associated begomovirus particles. To our knowledge, this is the first report from Bangladesh for identification and molecular characterization of MYVMD associated begomovirus complexes in kenaf. Several sporadic investigations were carried out to detect, isolate and characterize the causing viral agents of MYVMD in India [3,4,5,6,7,8,27,28,29]. However, accumulated molecular variability analysis of MYVMVs and associated betasatellites considering various evolutionary parameters are yet to be studied. Hence, with an aim to address this issue, we reported and discussed the sources of genetic variations, population dynamics and different forces acting on the evolution of MYVMVs and betasatellites. For theses analyses, we considered all the available MYVMV and betasatellite isolates so far (isolated from mesta plants or associated with mesta plants) in GenBank including the new isolates from Bangladesh (from this study). Moreover, we determined the potential linkage between the recombination events and presence of cSSR motifs in the studied begomovirus complexes. In a practical context, our findings can be useful to future epidemiological and evolutionary trend analyses towards development of comprehensive disease control strategies for begomoviruses.

## Materials and Methods

### Sample Collection and Viral Sequences

Naturally infected kenaf leaf samples showing typical yellow mosaic symptoms were collected from 7 kenaf producing districts of Bangladesh namely Faridpur, Gazipur, Jashore, Kishoreganj, Manikganj, Rangpur and Tangail. At least three samples were collected from each location. All the infected leaf samples were transported in an ice box to the laboratory and preserved at −80°C freezer until DNA isolation. The genomic DNA was isolated following the method described by Ghosh et al. 2009 [30] with necessary modifications. The complete genomic DNA of begomoviruses were PCR (Polymerase Chain Reaction) amplified with a set of forward (5’-GAGGGTGAGCTCTCCGAAGGTTGTAG-3’) and reverse (5’-AATTATGAGCTCTGGGAG-TGCGCAAG-3’) primers using DreamTaq DNA Polymerase (Thermo Fisher Scientific, USA). The isolated genomic DNAs were also subjected to amplify full length betasatellites using forward (5’-AGTAAGGGTACCACTACGCTACGCAG-3’) and reverse (5’-ATTAATGGTACCTACC-CTCCCAGGGGTAC-3’) primer pair. Both the amplified begomovirus genomes and betasatellites were individually cloned into the pGEM^®^-T Easy Vector Systems (Promega Corporation, USA). The colonies with desired clones were selected based on restriction patterns and three clones per isolate were sequenced bi-directionally for both begomovirus genomes and betasatellites using sanger sequencing facilities at Basic and Applied Research on Jute Project, Manik Mia Avenue, Dhaka, Bangladesh. The raw sequencing reads as chromatographs were assembled and edited with Geneious Prime software program.

### Phylogenetic Analysis, Recombination and Microsatellite Detection

In addition to the begomovirus and betasatellite sequences reported in this study, 28 MYVMV (23 from India and 5 from Pakistan) and 38 betasatellite (28 from India and 10 from Pakistan) sequences were downloaded from the GenBank and the details are presented in Table S1. All the sequences of downloaded MYVMVs and betasatellites were either isolated from or associated with mesta plants. MUSCLE alignment program was used to align the sequences [31] and Sequence Demarcation Tool (SDT v1.2) [32] was utilized to obtain the pairwise identities between the sequences. MUSCLE generated alignment for both the sequences were used to generate phylogenetic trees in molecular evolutionary genetics analysis version X (MEGA-X) [33] using a maximum-likelihood method with 1000 bootstrap replicates. Possible recombination events were detected with seven methods namely RDP, GENECONV, Bootscan, MaxChi, Chimaera, SiScan, and 3Seq using RDP4 (Recombination Detection Program version 4) [34]. Recombination predictions were only considered when the events were confirmed by at least 4 out of the 7 methods. Imperfect Microsatellite Extractor Version 2.1 (G-IMEx) was used to identify the presence of microsatellite in the viral sequences along with position [35] using the ‘Advance-Mode’ parameters as described in Chen et al. 2012 [36].

### Population Demography Analysis

Basic parameters of genetic variability and sequence diversity such as the number of variable sites (s), total number of mutations (η), nucleotide diversity per site (π), average number of nucleotide differences (k), the number of haplotype (h) and haplotype diversity (Hd) were calculated using DnaSP v6 [37]. To explore the demographic history of MYVMVs and associated betasatellies, several neutrality tests incorporated in DnaSP v6 such as Tajima’s D, Fu and Li’s D* and Fu and Li’s F* were implemented. The Tajima’s D test is based on the differences between the number of segregating sites and the average number of nucleotide differences whereas the Fu and Li’s D* and Fu and Li’s F* tests were based on the differences between the number of singletons with the total number of mutations and the average number of nucleotide differences between pairs of sequences respectively. Both the complete genome sequences and gene sequences (coat protein and rep protein of MYVMV and βC1 protein of betasatellite) were considered for all these analyses.

### Estimation of Selection Pressure

The acting evolutionary constraints or selection pressure on the coding regions of the viral populations were calculated based on the ratio of the nonsynonymous substitution rate (*d_N_*) to the synonymous substitution rate (*d_S_*). The nucleotide sequences of coat protein and rep protein of MYVMV and βC1 protein of betasatellite were used to calculate *d_N_/d_S_* ratio using single-likelihood ancestor counting (SLAC) method available in the DataMonkey server [38]. Next, a series of methods such as fixed-effects likelihood (FEL), SLAC and fast unbiased Bayesian approximation (FUBAR) incorporated in the DataMonkey server were utilized to find the sites under selection. The only positively selected sites identified by at least two of these methods were considered as under diversifying selection.

### Population Genetic Differentiation and Gene Flow Analysis

Permutation-based statistical tests Ks* and Z* [39] were considered to evaluate the extent of genetic differentiation among the viral populations. For measuring gene flow between the populations tested, F_ST_ and Nm values were taken into consideration. F_ST_ is calculated based on the mean number of differences between sequences sampled from the same subpopulation and the mean number of differences between sequences sampled from the two different subpopulations. Nm is based on the estimate of average divergence time of pairs of genes sampled from within a subpopulation and the average divergence time of genes sampled from different subpopulations [40]. All of these tests were calculated in DnaSP v6 considering the complete genome sequences and gene sequences of coat protein, rep protein and βC1 protein.

### Estimation of Nucleotide Substitution Rates

The rates of nucleotide substitution per site were estimated with the Bayesian Markov Chain Monte Carlo (MCMC) method available in the BEAST 1.10.4 package [41]. The nucleotide sequences of coat protein, rep protein and βC1 protein were used as input for BEAST analysis. For all the data set, both strict and relaxed (uncorrelated lognormal) molecular clocks were utilized along with the demographic model of piece-wise Bayesian skyline plot. Moreover, two substitution models (HKY and GTR) were used to generate BEAST files for each clock options (strict and relaxed). The resultant log files from BEAST analysis were analyzed into Tracer v1.6 to perform model comparison analyses (AICM analysis) and the best-fitting models were determined according to the AICM values. To ensure convergence, sufficient length of MCMC chains were run where an initial 10 % of the MCMC chains discarded as burn-in. The rates of nucleotide substitution per site were estimated from the best-fitting models of each data set using Tracer v1.6 program.

## Results

### Viral Sequence Analysis

Complete monopartite begomovirus genomes were PCR amplified with MYVMV specific primer pair from all the infected kenaf leaf samples collected from 7 districts of Bangladesh. However, betasatellite specific primer pairs were able to detect betasatellite genome in all the samples except the leaves collected from Rangpur district. All of these 13 amplified PCR products were cloned and sequenced for further analyses. The assembled contigs were pairwise aligned with known begomovirus and betasatellite sequences. The assembled monopartite genome sequences shared maximum nucleotide identities of 98.80 to 99.20% with MYVMVs, GenBank accession number NC_009088 and EF373060. Therefore, all the isolates of begomovirus reported in this study are identified as MYVMV isolates according to the demarcation threshold of begomovirus species [42]. On the other hand, the begomovirus associated betasatellite sequences shared maximum nucleotide identities of 96.90 to 98.30% with MYVMV associated betasatellites, GenBank accession number NC_009903 and DQ298137. As a result, all the isolates of begomovirus associated betasatellites reported in this study are identified as MYVMV associated betasatellite isolates according to the ICTV guidelines for betasatellite demarcation [43]. The pairwise comparison analysis of MYVMVs and betasatellites were presented in Figure 1a and Figure 1c respectively. The total 13 sequences reported in this study were deposited, assigned with accession number (MZ448274-MZ448286) and released in the GenBank. The details of this isolates with accession numbers are provided in Table S2. In addition to 7 MYVMVs and 6 associated betasatellites reported in this study, 28 MYVMV and 38 betasatellite sequences were downloaded from the GenBank and considered for further analysis.

**Figure 1.**
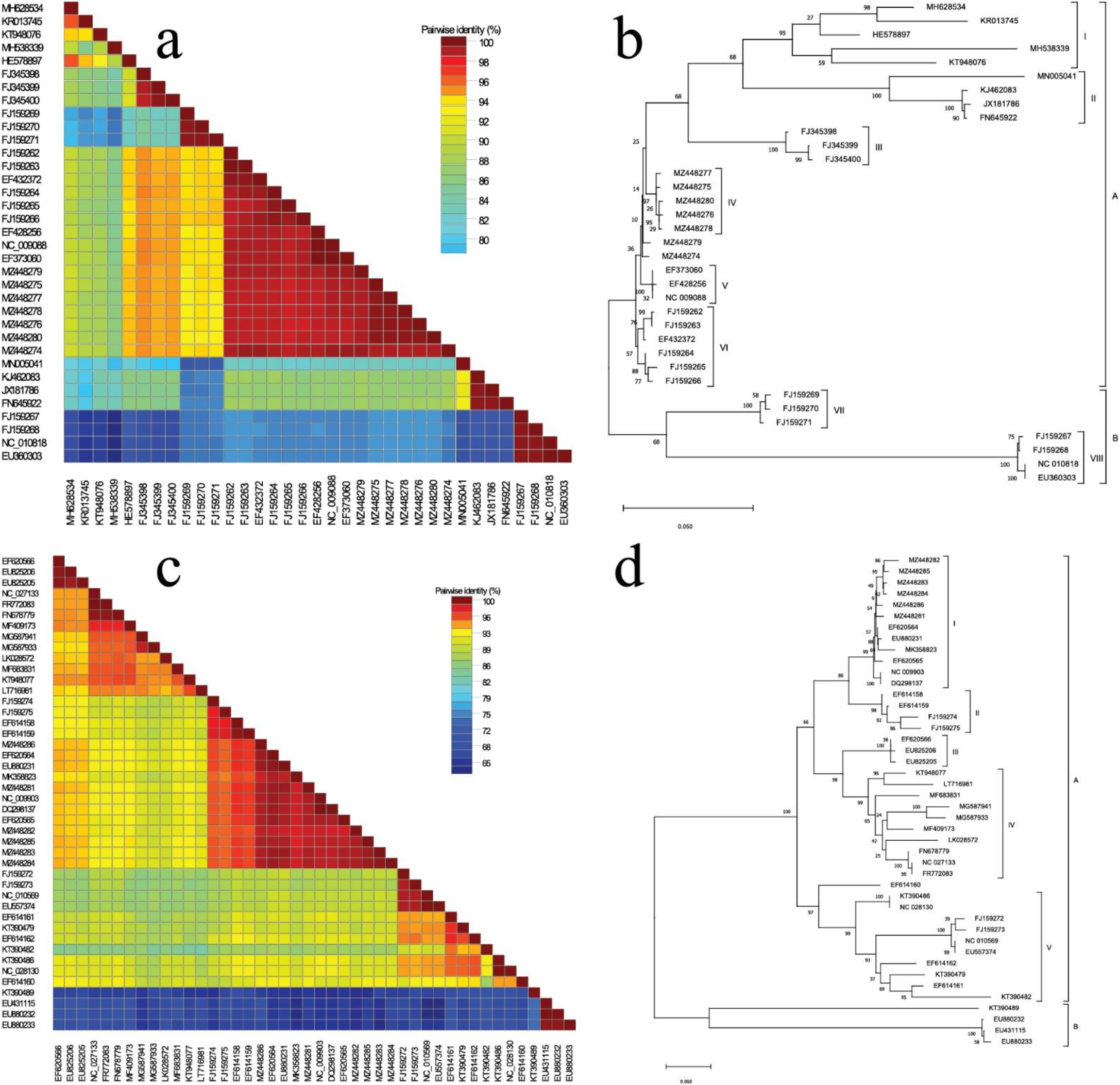
The pairwise comparison analysis in SDT v1.2 and phylogenetic analysis in MEGA-X using whole genome sequences of MYVMVs and betasatellites. Pairwise sequence identities (%) between the viruses are represented in different color codes. Each colored key represents a percentage to the identity score between two sequences. The trees were constructed using maximum-likelihood method with 1000 bootstrap replicates. **a**, Pairwise identity matrix of MYVMV isolates. **b**, Phylogenetic tree of MYVMV isolates. **c**, Pairwise identity matrix of betasatellite isolates. **d**, Phylogenetic tree of betasatellite isolates.

### Phylogenetic Relationship

As a part of the investigation of the genetic basis of evolution in MYVMV and associated betasatellites, we studied overall sequence variations and generated phylogenetic trees. MYVMV tree (Figure 1b) produced two major branches. All the MYVMV isolates from Pakistan were clustered into clade I under branch A while the Indian isolates were grouped into both the branches. Branch B was solely consisted of Indian isolates and all the Bangladeshi isolates are gathered under the branch A. Five of the 7 Bangladeshi isolates formed an independent clade however, rest two did not fall under any specific group. The phylogenetic tree constructed based on betasatellites was also divided into two major branches (Figure1d). Branch A was consisted of Bangladeshi, Indian and Pakistani isolates whereas only Indian isolates occupied the branch B. As like MYVMV tree, all the betasatellites from Pakistan were clustered under single clade, IV while betasatellites isolated from Bangladesh were separated into clade I with some Indian isolates.

### Recombination Analysis

Detection of recombination events and putative parental sequences were carried out with all the studied isolates of MYVMV and betasatellite. A total of 15 unique recombination events were detected in MYVMV isolates by at least 4 number of methods (Table 1). The analysis detected 20 isolates out of 35 as recombinants and among them, 12 isolates were found to have multiple events of recombination (Table 1 and Table S3). Maximum 4 events were possessed by each of the 3 South Indian isolates FJ159269, FJ159270 and FJ159271. All the isolates from Pakistan were detected as recombinants where 3 of them possessed one and rest 2 were contained with two recombination events. No isolate from Bangladesh was found as recombinants however, 3 of them (MZ448274, MZ448277 & MZ448280) were recognized as the parental sequences of both Indian and Pakistani recombinants (Table 1 and Table S3). Both inter-species and intra-species recombination events were detected and one of the parental sequences was found unknown in case of 8 events (Table 1).

**Table 1.**
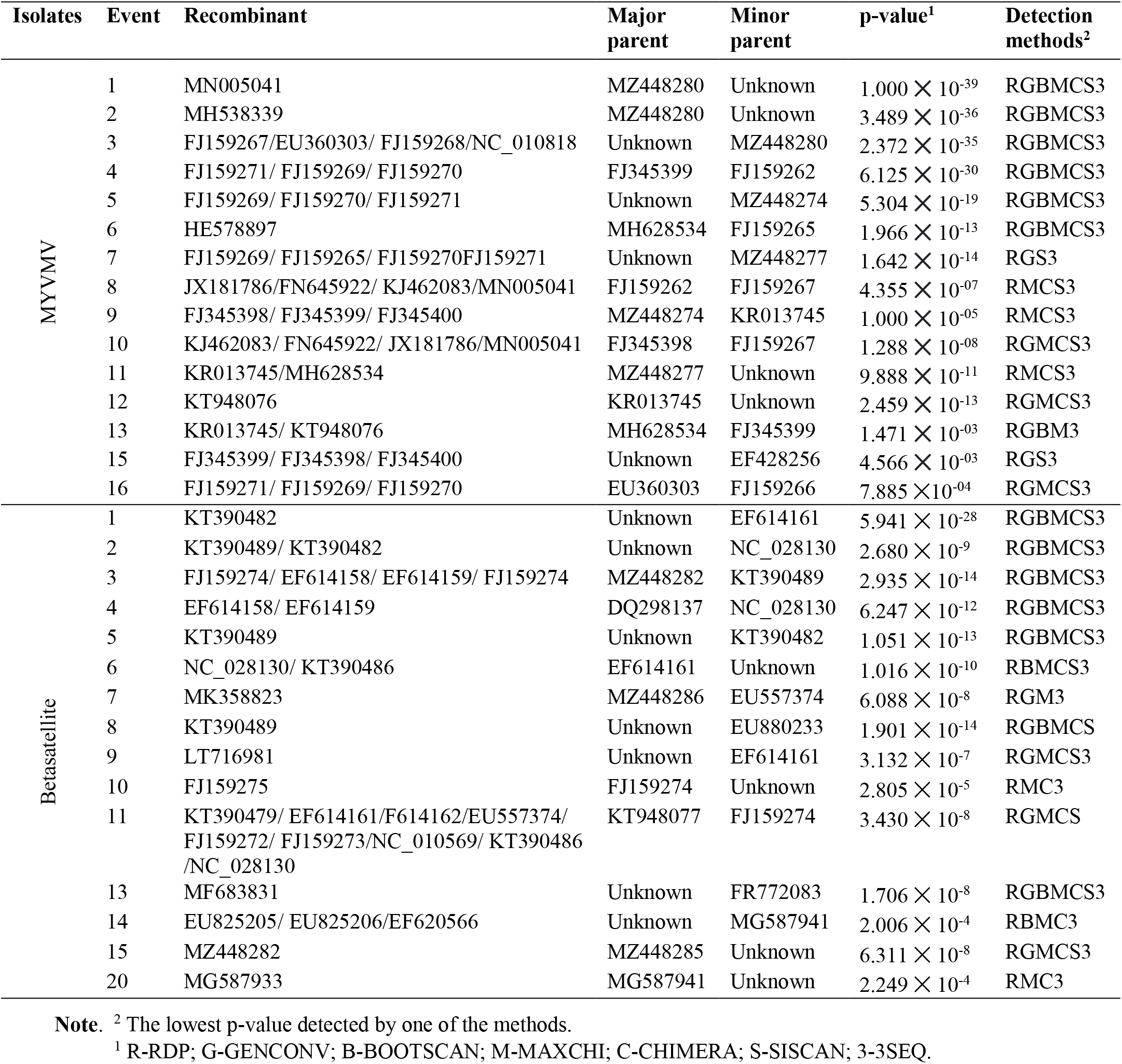
Recombination analyses of MYVMVs and associated betasatellites

In betasatellites, 23 out of 44 isolates were found to be recombinants possessing 15 unique recombination events (Table 1 and Table S4). Multiple events of recombination were contained by 7 isolates and Indian isolate KT390489 possessed 3 recombinantion events which is the highest among the betasatellites (Table S4). MZ448282 is the only Bangladeshi isolate found to be recombinant having another Bangladeshi isolate (MZ448285) as the major parent. The number of non-recombinant isolates was 9 and 7 out of 28 Indian and 10 Pakistani isolates respectively. The recombination events were found inter-species for 10 events and intra-species for 5 events. However, the detection program was unable to precisely determine the putative major parents in 7 recombination events and the number was 4 in case of minor parents.

### Association Between cSSR Motifs and Recombination Events

To determine the presence and position of cSSR motifs in the viral genomes, we performed sequence analysis using G-IMEx. The analysis detected cSSR motifs in all the genomes except one betasatellite isolate MG587933 from Pakistan. The number of cSSRs present per genome ranged from 1-4 for MYVMV and 1-3 for betasatellites (Figure S1a and Table S4). Two cSSRs were present in more than 60% isolates of both of the genomes and 4 cSSRs was only detected in KT390482, betasatellite from Uttar Pradesh, India. Most abundant types of cSSRs are (CG)3-x1-(T)7 & (AT)3-x-1-(T)7 in MYVMVs while (GA)3-x3-(TTG)4-x-2-(TG)3 and (A)9-x1-(A)6 are the prevailing types in betasatellites (Figure S1b).

Next, the association between the presence of recombination events and the cSSR motifs were studied. To observe the possible link of their co-existence, we counted the isolates that possessed both the cSSRs and recombination events, any of them or none of them. We analyzed that the presence of both the recombination event and cSSR were noticed in 57.14% of MYVMVs and 50% of betasatellites (Figure 2a). On contrary, 42.86% MYVMVs and 47.73% betasatellites were found to have cSSR motifs lacking recombination events in their genomes. However, this case is not found that both the cSSR and recombination event are missing. Moreover, we attempted to detect the trend of this relationship in the putative parental sequences identified by recombination detection program. Results showed that 61.54% parental sequence of MYVMVs and 47.73% of betasatellites were consisted of both the cSSRs and recombination events (Figure 2b) whereas 38.46% and 50% parental sequences were possessed cSSRs without any recombination event in MYVMVs and betasatellites respectively. Nevertheless, no recombination events were detected in the isolates lacking cSSR motifs.

**Figure 2.**
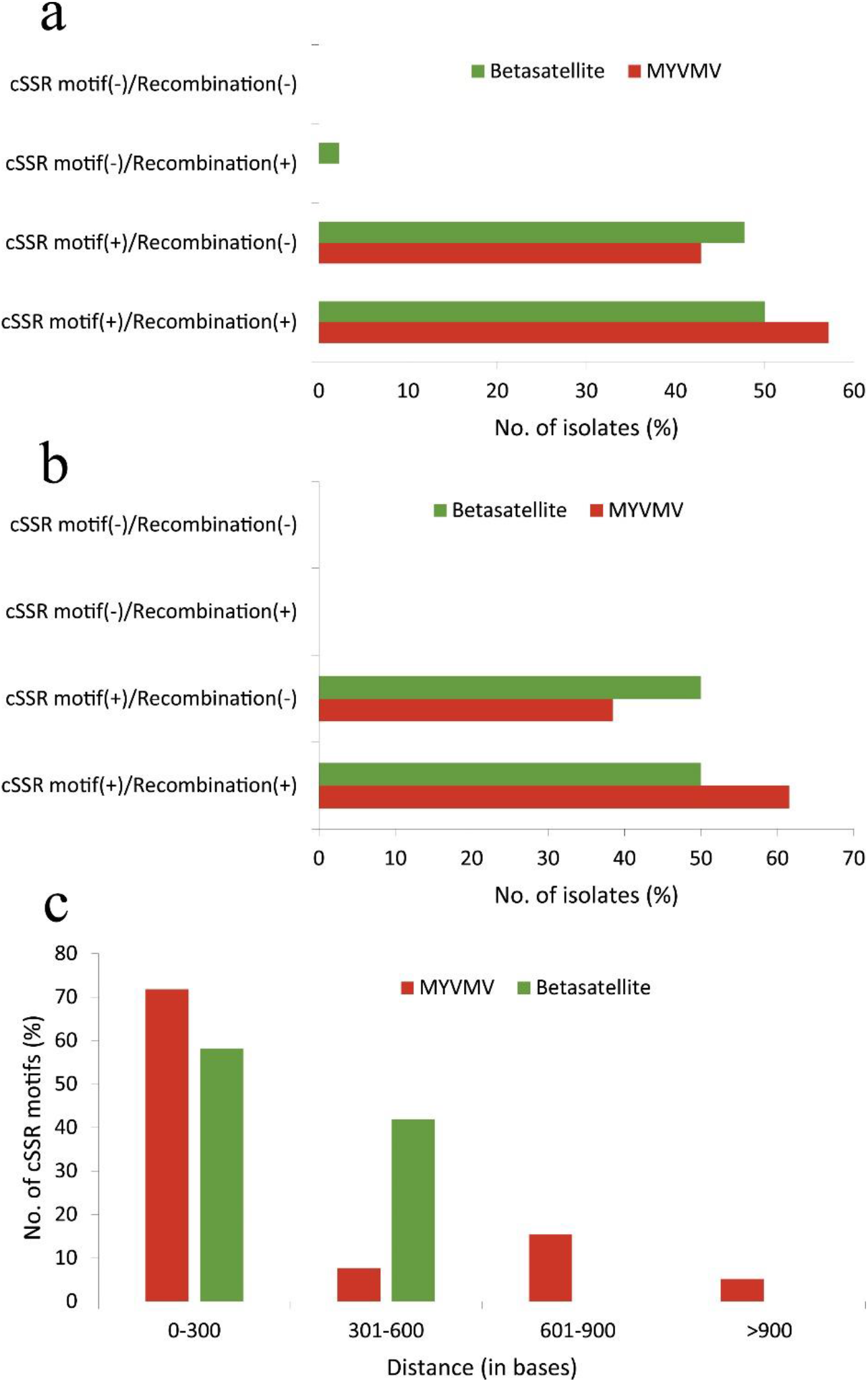
Association between recombination events and microsatellites detected in MYVMVs and betasatellites. **a,** The correlation between the cSSRs and recombination events detected in the viral genomes. ‘+’ and ‘−’ sign in the parenthesis indicate the presence or absence of cSSRs and recombination events detected in viral genomes. **b,** The correlation between the cSSRs and recombination events in the putative parental genomes identified in recombination detection software RDP4. ‘+’ and ‘−’ sign in the parenthesis indicate the presence or absence of cSSRs and recombination events detected in viral genomes. **c,** Physical proximity of the cSSRs and recombination breakpoints. Distances were calculated between each cSSR and the nearest recombination breakpoint in viral isolates which possess both cSSRs and recombination events.

Further, to find the location of cSSR motifs and recombination events we mapped their position onto the linearized genomes of MYVMV and betasatellite. The results indicated that, the recombination break points were distributed throughout the length of the MYVMV and betasatellite genomes compared to the position of cSSR motifs, although the majority of the break points in betasatellites were gathered in the satellite conserved region (SCR) (Figure 3a and Figure 3b). The cSSRs were found to be mostly located in the N-terminal rep protein and intergenic region, and non-coding and A-rich region of MYVMVs and betasatellites respectively. Furthermore, we sought to explore the correlation between cSSR motifs and recombination events in terms of their physical proximity which was calculated by measuring the nucleotide distance between each cSSR and the nearest recombination breakpoint if both were present in respective genome. Interestingly the majority of the cSSR motifs were found to be located within 300 bases of the nearest recombination break points in both MYVMVs and betasatellites (Figure 2c).

**Figure 3.**
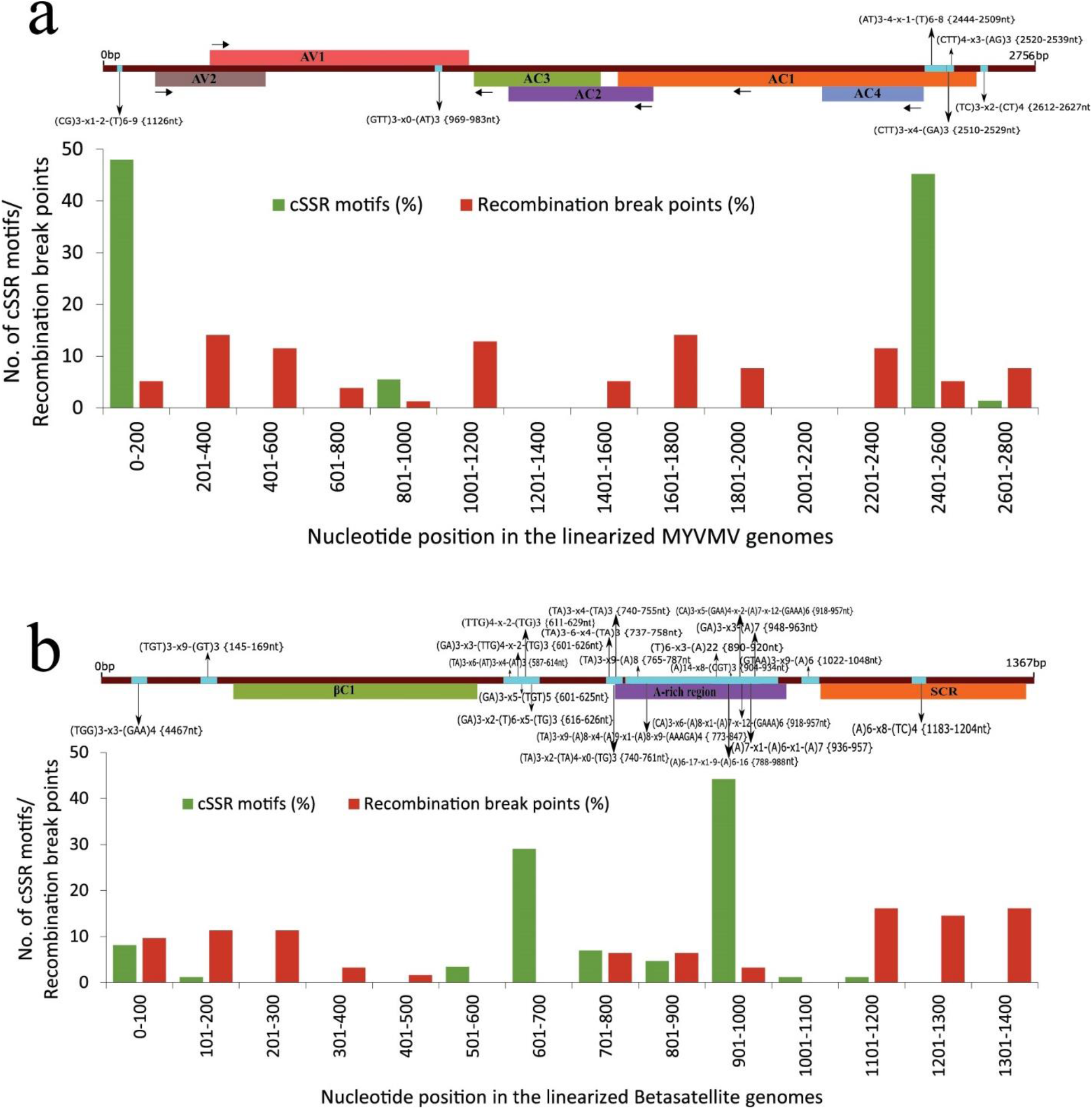
Genome-wide distribution and location of cSSR motifs and recombination breakpoints in MYVMV and betasatellite genomes. Schematic representation of a linearized MYVMV and betasatellite genome is placed over the respective graphical representation with similar scaling. In Schematic representation horizontal bars represents the genes, A-rich regions and conserved regions where applied. Horizontal arrow represents the directions of respective segment while vertical arrow represents the position of cSSR motifs. **a,** The distribution of total cSSR motifs and recombination breakpoints detected in the MYVMVs with schematic representation. AV1, coat protein; AV2, pre-coat protein; AC1, replication initiation protein or rep protein; AC2, transcriptional activator protein; AC3, replication enhancer protein and AC4, AC4 protein. **b,** The distribution of total cSSR motifs and recombination breakpoints detected in the betasatellites with schematic representation. βC1, βC1protein and SCR, satellite conserved region.

### Genetic Diversity and Population Dynamics

Nucleotide diversity was found high both in MYVMVs (π = 0.09) and betasatellites (π = 0.15) detected in DnaSP v6 software program (Table 2). The number of haplotypes were detected 33 & 40 along with 0.997 & 0.996 haplotype diversity in MYVMV and betasatellite genomes respectively. When considered geographically, the genetic variability was highest (π = 0.10 for MYVMVs and π = 0.18 for betasatellites) in Indian isolates. Pakistani isolates also possessed higher degree of variations whereas, very less nucleotide diversity (π = 0.006 for MYVMV and π = 0.017 for betasatellites) was observed in the isolates of Bangladesh. We also measured the genetic diversity in the genes coded for coat & rep protein of MYVMVs and βC1 protein of betasatellites. Similar trends of nucleotide diversity were found in gene analysis compared to whole genome analysis i.e. Indian isolates scored highest value of π while lowest value was achieved by Bangladeshi isolates (Table 2). The haplotypes number was found higher in coat protein compared to rep protein and the haplotype diversity was found 0.992, 1.000 & 0.976 for coat, rep and βC1 protein respectively.

**Table 2.**
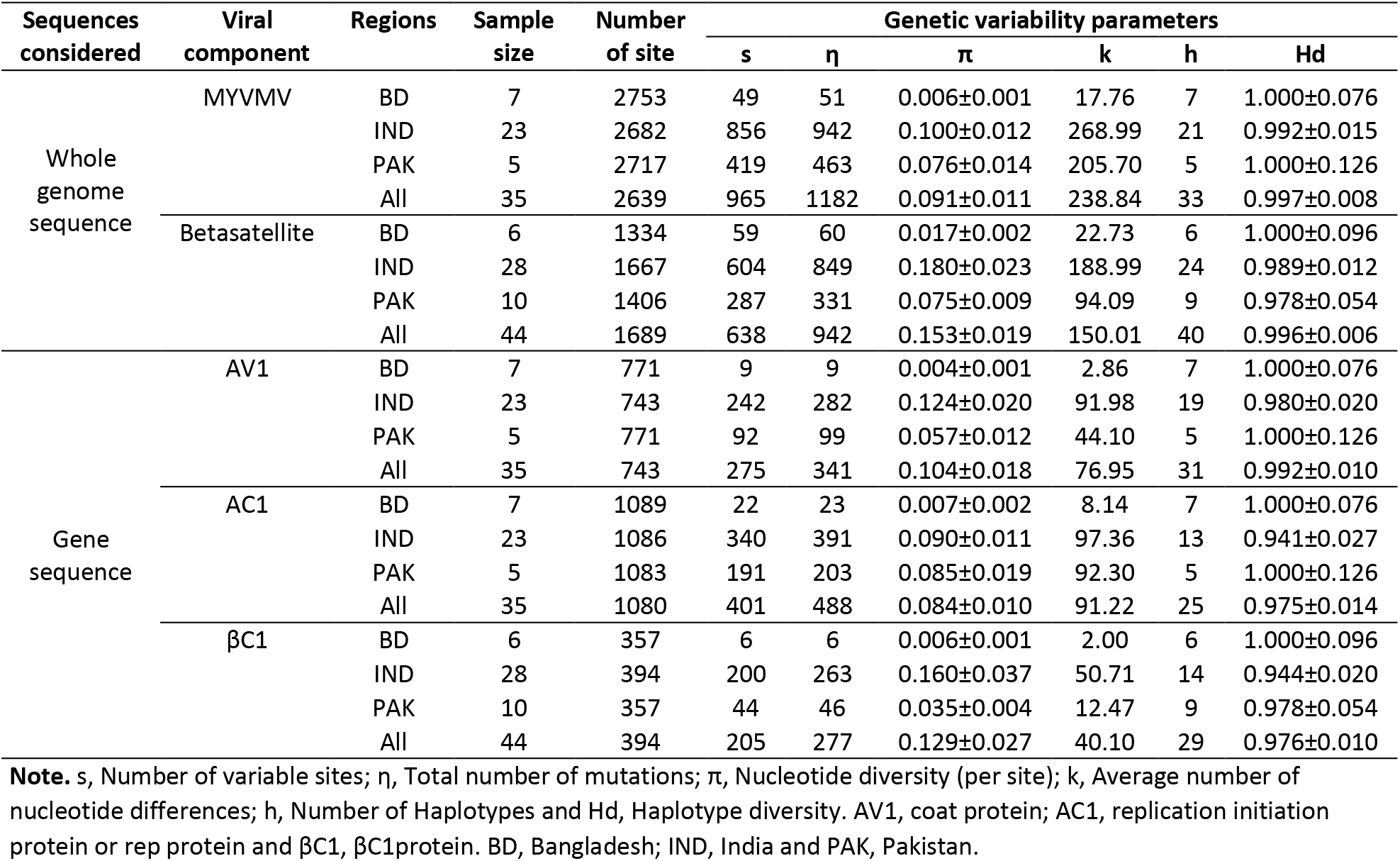
Genetic diversity of MYVMV and associated betasatellite isolates in different populations

To test the hypothesis of neutral selection between lineages and populations of MYVMV and betasatellite, several neutrality tests such as Tajima’s D, Fu and Li’s D* and Fu and Li’s F* were considered. Evolutionary events such as population expansion, bottlenecks and selection are speculated by Tajima’s D while Fu and Li’s D* tests are associated with population demographic expansion. All the values of these tests were found negative except the Fu and Li’s D* values in MYVMV populations when considered the complete genome sequences (Table 3). However, in country wise consideration, all the three values were calculated positive in MYVMV populations from India. Whether cumulative or country wise analyses were done, all the betasatellites produced negative values in all three indices in case of both whole genome and βC1 gene sequences (Table 3). The Fu and Li’s D* and Fu and Li’s F* values for coat and rep protein genes were estimated positive for Indian MYVMV populations in which values for coat protein genes were found statistically significant.

**Table 3.** Nutrality tests of MYVMV and associated betasatellite isolates in different populations

| Sequences considered | Viral component | Regions | Sample size | Tajima's D | Fu and Li's D* | Fu and Li's F* |
| --- | --- | --- | --- | --- | --- | --- |
| Whole genome sequence | MYVMV | BD | 7 | -0.84865 | -0.67052 | -0.78647 |
|  |  | IND | 23 | 0.22007 | 0.97331 | 0.86378 |
|  |  | PAK | 5 | -0.56878 | -0.36589 | -0.44331 |
|  |  | All | 35 | -0.64225 | 0.30508 | -0.03319 |
|  | Betasatellite | BD | 6 | -0.86664 | -0.80082 | -0.89404 |
|  |  | IND | 28 | -0.67317 | -0.20628 | -0.42727 |
|  |  | PAK | 10 | -0.97988 | -0.81145 | -0.96531 |
|  |  | All | 44 | -1.14060 | -0.53817 | -0.91052 |
| Gene sequence | AV1 | BD | 7 | -1.18108 | -1.32889 | -1.41888 |
|  |  | IND | 23 | 0.82744 | 1.67898** | 1.65640* |
|  |  | PAK | 5 | -0.54691 | -0.40299 | -0.46942 |
|  |  | All | 35 | -0.26901 | 0.91907 | 0.59838 |
|  | AC1 | BD | 7 | -0.74918 | -0.60431 | -0.70261 |
|  |  | IND | 23 | -0.32941 | 0.33409 | 0.14666 |
|  |  | PAK | 5 | -0.40234 | -0.28493 | -0.33599 |
|  |  | All | 35 | -0.87793 | -0.20287 | -0.52280 |
| | βC1 | BD | 6 | -1.36732 | -1.39992 | -1.48974 |
|  |  | IND | 28 | -0.97946 | -0.38526 | -0.68764 |
|  |  | PAK | 10 | -1.13759 | -1.25278 | -1.38224 |
|  |  | All | 44 | -1.36372 | -0.55617 | -1.02007 |

**Note.** AV1, coat protein; AC1, replication initiation protein or rep protein and βC1, βC1protein. BD, Bangladesh; IND, India and PAK, Pakistan; BD, Bangladesh; IND, India and PAK, Pakistan. \* $p < 0.05$ , \*\* $p < 0.02$

### Population Genetic Differentiation and Gene Flow

Independent statistical tests of population differentiation Ks* and Z* were performed using DnaSP v6 between geographical origins of isolates (Bangladesh, India and Pakistan) of MYVMV and betasatellite. The Ks* and Z* values were comparatively higher when analyzing the Indian isolates with Bangladeshi or Pakistani isolates while comparatively lower values were calculated when comparing Bangladeshi with Pakistani isolates (Table 4). The completely similar tendency was found whether full genomes or gene sequences were considered during the analyses. To find out the evidence of gene flow between those populations the number of effective migrants (Nm) values were calculated along with another genetic differentiation parameter F_ST_. Considering both genomes and gene sequences, the F_ST_ values ranged from 0.18603 to 0.54216 and 0.20264 to 0.73537 for MYVMV and betasatellite isolates respectively (Table 4). Regarding gene flow, the number of effective migrants (Nm) values were calculated <1 for all the data set except the AC1 gene sequences when comparing isolates of Bangladesh with Indian populations. Taking all the value trends under consideration, higher genetic differentiation and lower genetic flow were noticed between the isolates of Bangladesh and Pakistan.

**Table 4.** Genetic differentiation and gene flow analyses of MYVMV and associated betasatellite isolates in different populations

| Viral Component | Population | Genetic differentiation |  | Gene flow |  |
| --- | --- | --- | --- | --- | --- |
|  |  | Ks* | Z* | F<sub>ST</sub> | Nm |
| MYVMV genome sequence | BD-IND | 4.66688 | 4.87185 | 0.21595 | 0.91 |
|  | BD-PAK | 3.68327 | 2.70111 | 0.54216 | 0.21 |
|  | IND-PAK | 5.11954 | 4.77762 | 0.29490 | 0.60 |
| AV1 Gene sequence | BD-IND | 3.35434 | 4.83831 | 0.24454 | 0.77 |
|  | BD-PAK | 2.19763 | 2.70969 | 0.53150 | 0.22 |
|  | IND-PAK | 3.82111 | 4.81786 | 0.26398 | 0.70 |
| AC1 Gene sequence | BD-IND | 3.63016 | 4.94258 | 0.18603 | 1.09 |
|  | BD-PAK | 2.83008 | 2.72351 | 0.52229 | 0.23 |
|  | IND-PAK | 4.08841 | 4.76933 | 0.26358 | 0.70 |
| Betasatellite genome sequence | BD-IND | 4.64418 | 5.22571 | 0.25336 | 0.74 |
|  | BD-PAK | 3.92163 | 3.05172 | 0.64430 | 0.14 |
|  | IND-PAK | 4.65535 | 5.21305 | 0.26528 | 0.69 |
| βC1 Gene sequence | BD-IND | 3.05795 | 5.26652 | 0.20264 | 0.98 |
|  | BD-PAK | 2.00942 | 3.04262 | 0.73537 | 0.09 |
|  | IND-PAK | 3.13289 | 5.18609 | 0.27281 | 0.67 |

**Note.** AV1, coat protein; AC1, replication initiation protein or rep protein and βC1, βC1protein. BD, Bangladesh; IND, India and PAK, Pakistan.

### Selection Pressure Analysis

To detect the natural selection dominating on the coat, rep and βC1 protein, the ratio between the non-synonymous and synonymous substitution frequencies (*d_N_/d_S_*) were calculated in Datamonkey web server. The *d_N_/d_S_* ratio of coat protein was 0.131 which was approximately half compared to rep protein indicating coat protein under strong constrained for diversification (Table 5). The *d_N_/d_S_* ratio of the βC1 protein of betasatellites was calculated 0.266 which was slightly higher than rep protein and twice of coat protein of MYVMVs. Selection constraints operating on specific sites of the analyzed proteins were identified with three independent methods. General results reflected the *d_N_/d_S_* ratios (overall negative selection) showing a large number of sites under purifying selection than positive selection (Table 5). Only one site was found under positive selection in coat and rep protein by two methods whereas, no site was detected under diversifying selection by more than one method in βC1 protein. Not a single site was selected by SLAC method under positive selection however, the number of sites under negative selection detected by SLAC were comparatively lower than the sites detected by FEL and FUBAR (Table 5).

**Table 5.**
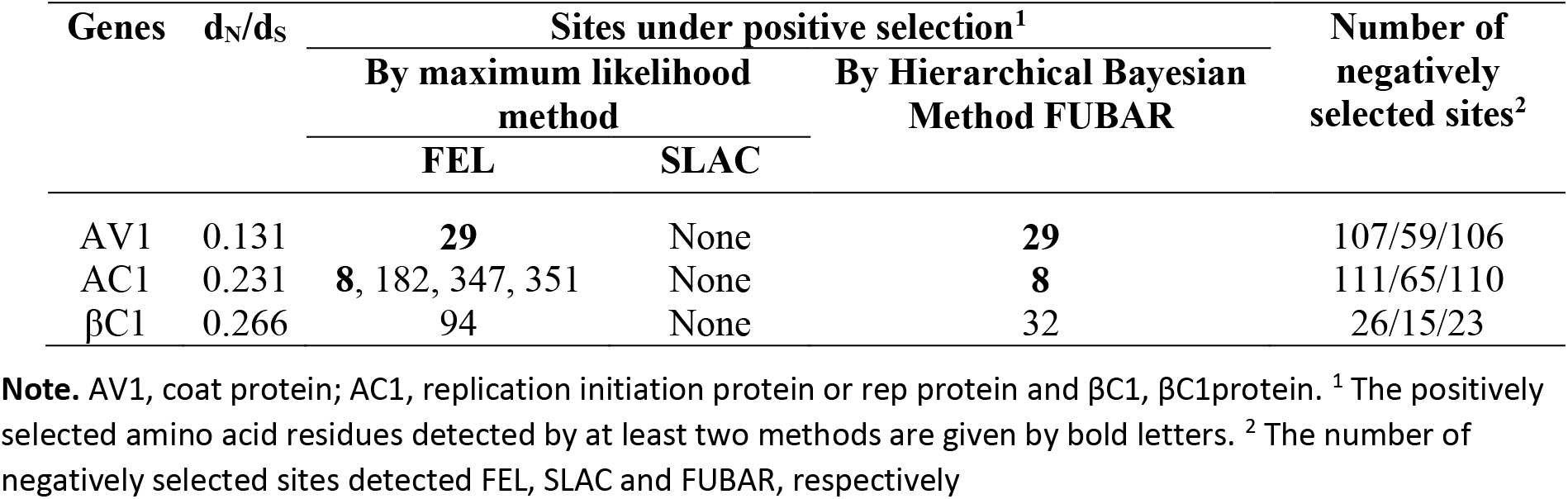
Selection pressure analyses in the selected genes of MYVMVs and associated betasatellites

### Estimation of Evolutionary Rates

Two substitution models HKY and GTR along with both strict and relaxed molecular clocks were used to detect nucleotide substitution rate nevertheless, GTR was selected as best-fitting model in model comparison for all the data sets. Relaxed molecular clock model was found better over strict clock in coat and rep protein data sets while strict molecular clock model was analyzed best-fitting in βC1 protein (Table 6). Notably, Bayesian skyline was used as demographic model for each data set and uncorrelated lognormal parameter was used for relaxed molecular clock model. The mean rate of substitution for coat protein was found 5.99 × 10^−3^ (95% HPD 1.34 × 10^−3^ to 1.11 × 10^−2^) nucleotide substitutions/site/year which is higher than the rep protein estimated mean rate of 2.59 × 10^−4^ with 95% HPD 4.18 × 10^−6^ to 6.68 × 10^−4^ nucleotide substitutions/site/year (Table 6). The βC1 protein has been estimated with 5.39 × 10^−6^ and 7.15 × 10^−23^ to 3.40 × 10^−5^ nucleotide substitutions/site/year as the mean rate and 95% HPD respectively pointing out the lower rate of substitution of betasatellite protein than two MYVMV proteins.

**Table 6.** Estimation of evolutionary rates in the selected genes of MYVMVs and associated betasatellites

| Parameters | Coat protein (AV1) | Rep protein (AC1) | βC1 protein (βC1) |
| --- | --- | --- | --- |
| Best-fit molecular clock model | Relaxed (lognormal) | Relaxed (lognormal) | Strict |
| Best-fit substitution model | GTR | GTR | GTR |
| No. of sequences | 35 | 35 | 44 |
| Sequence length (nt) | 771 | 1101 | 394 |
| Chain length (millions) | 80 | 60 | 100 |
| Mean substitution rate (nucleotides/site/year) | $5.99 \times 10^{-3}$ | $2.59 \times 10^{-4}$ | $5.39 \times 10^{-6}$ |
| HPD substitution rate (nucleotides/site/year) | $1.34 \times 10^{-3}$ to $1.11 \times 10^{-2}$ | $4.18 \times 10^{-6}$ to $6.68 \times 10^{-4}$ | $7.15 \times 10^{-23}$ to $3.40 \times 10^{-5}$ |

**Note.** HPD, 95% high probability density

## Discussion

Mesta plant, which has two cultivated species *Hibiscus cannabinus* (kenaf) and *Hibiscus sabdariffa* (roselle), beside its’ traditional role as the producer of natural fiber is a promising alternative source of quality pulp for tree-free paper manufacturing industries addressing the climate changing issues. The MYVMD, characterized with a typical yellow vein mosaic symptom, is one of the major factors for hindering the ultimate yield of mesta plants [3,4]. MYVMV and several betasatellites were proved to be the causal agents of the disease [6,7,8]. The mesta crops cultivated in Bangladesh are also affected by this mosaic disease seriously however, no study was reported to detect and characterize the causal agents of the disease in Bangladesh. In this circumstances, beside the isolation and identification of the causing agents of MYVMD from the infected samples of Bangladesh, we attempted to perform detailed molecular and evolutionary study in available MYVMVs and betasatellites associated with mesta plants including new isolates from Bangladesh.

The begomovirus particles isolated from 7 leaves samples of Bangladeshi kenaf plants were identified as MYVMV according to the demarcation threshold of begomovirus species [42] and the betasatellites isolated from 6 of these samples were identified as MYVMV associated betasatellites according to the ICTV guidelines for betasatellite demarcation [43]. The betasatellite specific primer did not able to amplify any product from the sample collected from Rangpur districts of Bangladesh. It was evident that in the absence of a betasatellite monopartite begomovirus could evolve [44]. Surprisingly, the MYVMV isolated from Rangpur district (MZ448279) was not clustered with other Bangladeshi isolates under clade IV in the phylogenetic analyses rather it was placed independently in the tree (Figure 1b). The isolates from eastern part of India were grouped in clade III and VI under branch A while MYVMVs from North and South India were separated into clade VII and VIII under major branch B. Notably, all the Pakistani isolates disregarding MYVMVs or betasatellites were set exclusively under single clade (Figure 1b and Figure 1d). However, isolates of Bangladesh were gathered under clade I with some Indian isolates in betasatellite tree. It can be speculated form the phylogenetic tree analyses that geographic location has had major influence in the evolution of MYVMVs and associated betasatellites. Although 3 Tomato leaf curl Joydebpur DNA betasatellites (EU880232, EU880233 and EU431115) along with Croton yellow vein mosaic betasatellite (KT390489) were isolated from *Hibiscus cannabinus,* they were distinctly separated in the betasatellite tree forming a major branch B (Figure 1d).

Recombination was proved to be a crucial factor in the evolution of viral species under Geminiviridae family [45,46]. In accordance, more than 50% MYVMV isolates were detected as recombinants in this study (Table 1 and Table S3). The majority of the recombination break points were located within the region of AV1/AV2 and AC1 genes (Figure 3a). The identification of AV1/AV2 and AC1 regions as the recombination hotspots is in agreement with some previous reports [20,21]. Recombination events 2,6,11 and12 were solely detected in Pakistani isolates while the rest of the maximum events were distributed among more than one Indian isolates (Table 1). Interestingly no Bangladeshi isolates were found as recombinant nevertheless, 3 of them (MZ448274, MZ448277 & MZ448280) were recognized as the parental sequences of both Indian and Pakistani recombinants suggesting the ancient and conservative lineages of MYVMV in Bangladesh. In case of betasatellites also, more than 50% of isolates were found to be recombinants possessing 15 unique recombination events (Table 1 and Table S4). Event 9,13 and 20 were solely shared by Pakistani isolates however, recombinant LT716981 was detected to have an Indian isolate as minor parent. Recombination event 11 was shared by 9 Indian isolates which was maximum per event and interestingly a Pakistani isolate KT948077 was identified as the major parent of this event (Table 1). The overall results indicated more diversified recombination in betasatellites compared to MYVMVs.

In recent time, the cSSR motifs were reported to be potentially linked with the recombination of begomoviruses [20,21]. Consequently, we detected the presence and position of cSSR motifs in the studied data sets and tried to figure out their relation with recombination breakpoints. The analyses indicated that almost 100% of isolates that were detected as recombinants also found to have cSSR motifs in their genomes (Figure 2a). Similar results were found when considered the putative parents of the recombination events in MYVMVs and betasatellites (Figure 2b). Moreover, the presence of recombination events in the absence of cSSR motifs in particular isolates is very rare which postulated the possible link between their co-existence and these observations are consistent with the findings of Kumar et al 2017. The genome wide distribution of cSSRs and recombination breakpoints of MYVMVs and betasatellites (Figure 3a and Figure 3b) depicted their appearance throughout the length of the genome however, cSSR motifs were mostly confined to N-terminal rep protein and intergenic region, and non-coding and A-rich region of MYVMVs and betasatellites respectively. In addition to the genome wide mapping, we calculated the physical proximity between each cSSR motif and the nearest recombination breakpoint and excitingly observed that 72% motifs in MYVMVs and 58% in betasatellites were located within 300 bases of corresponding break points (Figure 2c). All of these results are in well accordance with the previous studies of George et al. 2015 [21] and Kumar et al. 2015 [47] in begomoviruses. The possible reasons behind this association were speculated in some reports where these repetitive sequences were supposed to be an inducing agent triggering crossing-over or a source of signal for recombination [21,48,49] however, these ideas are yet to be validated experimentally.

Low genetic variability tends to be common in natural populations of plant viruses [50] however, in this study the overall genetic diversity was found very high in both whole genome and gene based study of MYVMVs and betasatellites with nucleotide diversity ranging from 0.084 to 0.153 (Table 2) when calculated for the whole population. These values were found much more higher compared to other begomoviruses [51,52]. In geographical consideration, the highest diversity was shown by Indian isolates ranging from 0.090 to 0.180 compared to Pakistan and Bangladesh. Bangladeshi isolates showed comparatively very low variability with nucleotide diversity ranging from 0.004 to 0.017. In congruence with the recombination analyses of this study, betasatellites were observed to maintain more genetic diversity among its’ isolates compared to MYVMVs and this finding is supported by the fact that higher genetic variability may be linked with more frequent and diversified recombination [51,52].

Three statistical tests, Tajima’s D, Fu and Li’s D* and Fu and Li’s F* suggested the molecular diversity patterns between lineages and populations. Apparently all the three values were found negative in the populations of Bangladesh and Pakistan (Table 3) indicating low frequency of polymorphism maintained in the populations of these two regions caused either by background selection, genetic hitchhiking or recent expansion. In contrast, Indian populations produced positive values in all three indices (only found negative Tajima’s D in rep protein) whether considered whole genome or gene sequences of MYVMVs. Moreover, the Fu and Li’s D* and Fu and Li’s F* values of coat protein were found much higher and statistically significant indicating polymorphism with high frequency which suggested that balancing selection or expansion has occurred in Indian populations. Two independent statistical tests Ks* and Z* values were calculated to measure population differentiation between three regions. In all the data sets, Ks* and Z* values were found lower in Bangladesh-Pakistan compared to Bangladesh-India and later one produced nearly similar values of India-Pakistan data sets suggesting high genetic differentiation between the populations of Bangladesh and Pakistan (Table 4). To estimate the degree of gene flow between geographical origins, the number of effective migrants (Nm) values were estimated along with F_ST_ which is a genetic differentiation index correlated with Nm. Generally, lower gene migration was observed in the populations with high degree of differentiation [53]. In accordance with the previous results based on Ks* and Z*, higher value of differentiation (F_ST_) was calculated with lower degree of gene flow (Nm) in all the Bangladesh-Pakistan data sets (Table 4). All these tests reinforce the finding that the populations of Bangladesh-Pakistan data set are distantly related with each other compared to Bangladesh-India and India-Pakistan. One possible explanation for this correlation could be the relatively far geographical distance between Bangladesh and Pakistan.

In this study, the *d_N_/d_S_* ratios of coat, and rep proteins of MYVMVs and βC1 protein of betasatellites were calculated <1 indicating a negative or purifying selection, albeit a different degree of selection pressure over each gene was observed (Table 5). The results suggested a stronger selection constraints operating on coat protein compared to rep protein which was exposed to comparatively weak selection pressure. In accordance with the overall purifying selections identified by *d_N_/d_S_* ratios, the number of individual sites under negative selection outnumbered the positively selected sites in all the data types and this result is supported by the earlier reports which emphasized the role of purifying selection on begomovirus evolution [14,46,52]. Notably, for having structural constraints and maintaining specific molecular interactions, many plant viruses were found to have purifying selection as the main evolutionary force [50]. No site under diversifying selection was detected by SLAC which is known to be a more conservative method to detect sites under selection [54]. The amino acid 8, only positively selected site in rep protein (Table 5), was part of the N-terminal region comprising of different conserved motifs required for virus replication [55]. The estimated nucleotide substitution rates were found very high in coat and rep protein compared to βC1 protein (Table 6). The observation of high substitution rate in begomovirus coat protein is consistent with some other studies [14,15,20]. We found the strongest purifying selection pressure (*d_N_/d_S_* = 0.131) in coat protein and its’ high nucleotide substitution rate is very likely to result from this strong negative selection acting on coat protein. The possible reasons for achieving such a high nucleotide substitution rate by begomoviruses might include the usage of error prone DNA polymerases in virus replication [14,56] or spontaneous mutations.

## Conclusions

This study reports the MYVMVs and betasatellites associated with MYVMD of kenaf plants in Bangladesh along with their molecular characterization. To our knowledge, this is the first report from Bangladesh on MYVMD associated begomovirus complex. This study also describes detailed molecular evolutionary analyses of available MYVMVs and associated betasatellites based on different parameters. The findings generated through our studies reinforces the fact that recombination plays an important role in the evolution and diversification of begomoviruses and betasatellites. A potential linkage is also demonstrated between the recombination events and presence of cSSR motifs. In addition to stating high degree of evolutionary rate and genetic variability found in MYVMVs and betasatellites, the study describes the relation between various index of population genetics addressing the geographic locations of the isolates. The findings of this study on different evolutionary forces that shape the emergence and diversification of MYVMVs and associated betasatellites may provide directions for future evolutionary study of different species of begomoviruses.

## Author Contributions

Rasel Ahmed conceived and designed the study, performed the experiments, performed bioinformatics analyses and wrote the manuscript. Rasel Ahmed, Borhan Ahmed, Md. Wali Ullah and Rajnee Hasan contributed to data analyses and graphics representation. Borhan Ahmed, Md. Wali Ullah and Rajnee Hasan contributed to critical reviewing and editing the manuscript. Borhan Ahmed and Md. Wali Ullah supervised the study.

