## Supplementary material for "Molecular evolution and genetic analysis of Mesta yellow vein mosaic virus and associated betasatellites": Figure S1

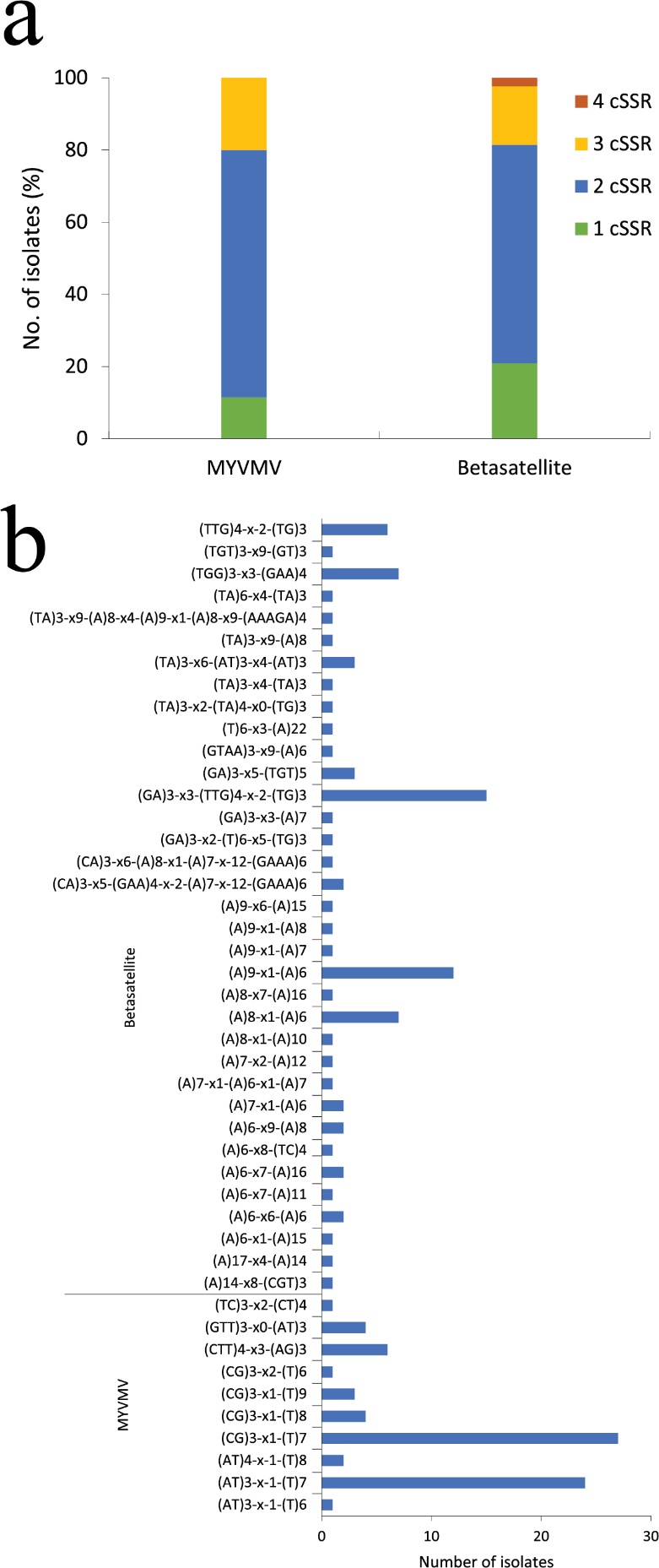

**Figure S1.** Detection of cSSR motifs in MYVMV and Betasatellite genomes. **a,** Distribution of the total number of cSSRs detected in the genomes. The digits (1–4) denote the number of microsatellites present in the given sequence. **b,** The frequency of cSSR motifs noticed among the genomes.
