## Supplementary material for "Molecular evolution and genetic analysis of Mesta yellow vein mosaic virus and associated betasatellites": Table S1-S4

**Table S1.** Sequences downloaded from GenBank

| **Accession Number** | **Length (bases)** | **Country:Region** |
| --- | --- | --- |
| **MYVMV sequences** | | |
| MN005041 | 2743 | India: Krishnanagar, West Bengal |
| MH628534 | 2749 | Pakistan |
| MH538339 | 2742 | Pakistan |
| KT948076 | 2756 | Pakistan: Lahore |
| KR013745 | 2750 | Pakistan |
| KJ462083 | 2742 | India: Varanasi, Uttar Pradesh |
| JX181786 | 2742 | India |
| HE578897 | 2752 | Pakistan:Lahore |
| FN645922 | 2742 | India:Haryana |
| FJ345398 | 2752 | India: East India, Barrackpore |
| FJ345399 | 2752 | India: East India, Haringhata |
| FJ345400 | 2752 | India: East India, Bongaon |
| FJ159262 | 2752 | India: East India, Bongaon |
| FJ159263 | 2752 | India: East India, Haringhata |
| FJ159264 | 2752 | India: East India, Raigunj |
| FJ159265 | 2752 | India: East India, Coochbehar |
| FJ159266 | 2752 | India: East India, Balurghat |
| FJ159267 | 2737 | India: North India, Kaisargunj |
| FJ159268 | 2737 | India: North India, Bhanga |
| FJ159269 | 2752 | India: South India, Amadalavalasa |
| FJ159270 | 2752 | India: South India, Vizianagaram |
| FJ159271 | 2752 | India: South India, Srikakulam |
| NC_010818 | 2737 | India: North India, Bahraich |
| EU360303 | 2737 | India: North India, Bahraich |
| EF432372 | 2752 | India |
| EF428256 | 2752 | India |
| NC_009088 | 2728 | India |
| EF373060 | 2728 | India |
| **Betasatellite sequences** | | |
| FJ159272 | 1362 | India: South India, Vizianagaram |
| FJ159273 | 1362 | India: South India, Srikakulam |
| FJ159274 | 1351 | India: East India, Raigunj |
| FJ159275 | 1351 | India: East India, Balurghat |
| NC_010569 | 1362 | India: South India, Amadalavalasa |
| EU557374 | 1362 | India: South India, Amadalavalasa |
| NC_009903 | 1354 | India |
| EF614158 | 1351 | India: East India, Bongaon |
| EF614159 | 1351 | India: East India, Haringhata |
| EF614160 | 1355 | India: North India, Bahraich |
| EF614161 | 1351 | India: North India, Bhangha |
| EF614162 | 1352 | India: North India, Kaisarganj |
| DQ298137 | 1354 | India |
| KT390489 | 1367 | India: Uttar Pradesh, Mirzapur |
| KT390486 | 1353 | India: Uttar Pradesh, Varanasi |
| KT390482 | 1365 | India: Uttar Pradesh, Mirzapur |
| KT390479 | 1359 | India: Uttar Pradesh, Varanasi |
| NC_028130 | 1353 | India: Uttar Pradesh, Varanasi |
| EF620566 | 1343 | India: North India, Bahraich |
| EU431115 | 1367 | India: Amadalavalasa, South India |
| EU825206 | 1343 | India: North India, Bhangha |
| EU825205 | 1343 | India: North India, Kaisarganj |
| EU880233 | 1367 | India: South India, Ponduru |
| EU880232 | 1367 | India: South India, Amadalavalasa |
| EF620564 | 1345 | India: East India, Barrackpore |
| EU880231 | 1345 | India: East India, Bongaon |
| EF620565 | 1347 | India: East India, Haringhata |
| MK358823 | 1345 | India: West Bengal,Siliguri |
| NC_027133 | 1346 | Pakistan:Faisalabad |
| FR772083 | 1346 | Pakistan:Faisalabad |
| FN678779 | 1346 | Pakistan:Lahore |
| LK028572 | 1352 | Pakistan |
| KT948077 | 1362 | Pakistan: Lahore |
| LT716981 | 1352 | Pakistan:Lahore |
| MG587941 | 1322 | Pakistan |
| MG587933 | 1309 | Pakistan |
| MF683831 | 1344 | Pakistan: Lahore |
| MF409173 | 1349 | Pakistan: Lahore |

**Table S2.** Sequences isolated from Bangladesh

| **Accession Number** | **Length (bases)** | **Country:Region** |
| --- | --- | --- |
| **MYVMV sequences** | | |
| MZ448274 | 2753 | Bangladesh:Faridpur |
| MZ448275 | 2754 | Bangladesh:Gazipur |
| MZ448276 | 2755 | Bangladesh:Jashore |
| MZ448277 | 2754 | Bangladesh:Kishoreganj |
| MZ448278 | 2754 | Bangladesh:Manikganj |
| MZ448279 | 2753 | Bangladesh:Rangpur |
| MZ448280 | 2754 | Bangladesh:Tangail |
| **Betasatellite sequences** | | |
| MZ448281 | 1347 | Bangladesh:Faridpur |
| MZ448282 | 1346 | Bangladesh:Gazipur |
| MZ448283 | 1347 | Bangladesh:Jashore |
| MZ448284 | 1345 | Bangladesh:Kishoreganj |
| MZ448285 | 1345 | Bangladesh:Manikganj |
| MZ448286 | 1348 | Bangladesh:Tangail |

**Table S3.** Identification of the recombination breakpoints and cSSR motifs in MYVMV isolates

| **Accession number** | **cSSR analysis** | | | **Recombination analysis** | | | | | | |
| --- | --- | --- | --- | --- | --- | --- | --- | --- | --- | --- |
|  | **cSSR motif** | **cSSR position** | | **Event number** | **Recombination breakpoints** | | **Putative parental sequences** | | **RDP methods^1^** | ***p*-value^2^** |
|  |  | **Start** | **End** |  | **Start** | **End** | **Major** | **Minor** |  |  |
| EF373060 | (CG)3-x1-(T)7 | 11 | 24 | - | - | - | - | - | - | - |
|  | (AT)4-x-1-(T)8 | 2444 | 2458 |  |  |  |  |  |  |  |
| EF428256 | (CG)3-x1-(T)7 | 11 | 24 | - | - | - | - | - | - | - |
|  | (AT)3-x-1-(T)7 | 2471 | 2482 |  |  |  |  |  |  |  |
| EF432372 | (CG)3-x1-(T)7 | 11 | 24 | - | - | - | - | - | - | - |
|  | (GTT)3-x0-(AT)3 | 969 | 983 |  |  |  |  |  |  |  |
|  | (AT)3-x-1-(T)7 | 2471 | 2482 |  |  |  |  |  |  |  |
| EU360303 | (CG)3-x1-(T)7 | 11 | 24 | 3 | 1751 | 31 | Unknown | MZ448280 | RGBMCS3 | 2.372X10-35 |
|  | (AT)3-x-1-(T)7 | 2457 | 2468 |  |  |  |  |  |  |  |
| FJ159262 | (CG)3-x1-(T)7 | 11 | 24 | - | - | - | - | - | - | - |
|  | (AT)3-x-1-(T)7 | 2471 | 2482 |  |  |  |  |  |  |  |
| FJ159263 | (CG)3-x1-(T)7 | 11 | 24 | - | - | - | - | - | - | - |
|  | (AT)3-x-1-(T)7 | 2471 | 2482 |  |  |  |  |  |  |  |
| FJ159264 | (CG)3-x1-(T)7 | 11 | 24 | - | - | - | - | - | - | - |
|  | (GTT)3-x0-(AT)3 | 969 | 983 |  |  |  |  |  |  |  |
|  | (AT)3-x-1-(T)7 | 2471 | 2482 |  |  |  |  |  |  |  |
| FJ159265 | (CG)3-x1-(T)7 | 11 | 24 | 7 | 297 | 466 | Unknown | MZ448277 | RGS3 | 1.642X10-14 |
|  | (GTT)3-x0-(AT)3 | 969 | 983 |  |  |  |  |  |  |  |
|  | (AT)3-x-1-(T)7 | 2471 | 2482 |  |  |  |  |  |  |  |
| FJ159266 | (CG)3-x1-(T)7 | 11 | 24 | - | - | - | - | - | - | - |
|  | (GTT)3-x0-(AT)3 | 969 | 983 |  |  |  |  |  |  |  |
|  | (AT)3-x-1-(T)7 | 2471 | 2482 |  |  |  |  |  |  |  |
| FJ159267 | (CG)3-x1-(T)7 | 11 | 24 | 3 | 1701 | 2715 | Unknown | MZ448280 | RGBMCS3 | 2.372X10-35 |
|  | (AT)3-x-1-(T)7 | 2457 | 2468 |  |  |  |  |  |  |  |
| FJ159268 | (CG)3-x1-(T)7 | 11 | 24 | 3 | 1751 | 31 | Unknown | MZ448280 | RGBMCS3 | 2.372X10-35 |
|  | (AT)3-x-1-(T)7 | 2457 | 2468 |  |  |  |  |  |  |  |
| FJ159269 | (CG)3-x1-(T)7 | 11 | 24 | 4 | 2204 | 296 | FJ345399 | FJ159262 | RGBMCS3 | 6.125X10-30 |
|  | (AT)3-x-1-(T)7 | 2471 | 2482 | 7 | 297 | 466 | Unknown | MZ448277 | RGS3 | 1.642X10-14 |
|  |  |  |  | 16 | 467 | 619 | EU360303 | FJ159266 | RGMCS3 | 7.885X10-04 |
|  |  |  |  | 5 | 1031 | 1895 | Unknown | MZ448274 | RGBMCS3 | 5.304X10-19 |
| FJ159270 | (CG)3-x1-(T)7 | 11 | 24 | 4 | 2204 | 296 | FJ345399 | FJ159262 | RGBMCS3 | 6.125X10-30 |
|  | (AT)3-x-1-(T)7 | 2471 | 2482 | 7 | 309 | 524 | Unknown | MZ448277 | RGS3 | 1.642X10-14 |
|  |  |  |  | 16 | 467 | 627 | EU360303 | FJ159266 | RGMCS3 | 7.885X10-04 |
|  |  |  |  | 5 | 1031 | 1895 | Unknown | MZ448274 | RGBMCS3 | 5.304X10-19 |
| FJ159271 | (CG)3-x1-(T)7 | 11 | 24 | 16 | 959 | 620 | EU360303 | FJ159266 | RGMCS3 | 7.885X10-04 |
|  | (AT)3-x-1-(T)7 | 2471 | 2482 | 4 | 2204 | 296 | FJ345399 | FJ159262 | RGBMCS3 | 6.125X10-30 |
|  |  |  |  | 7 | 309 | 524 | Unknown | MZ448277 | RGS3 | 1.642X10-14 |
|  |  |  |  | 5 | 1031 | 1895 | Unknown | MZ448274 | RGBMCS3 | 5.304X10-19 |
| FJ345398 | (CG)3-x1-(T)7 | 11 | 24 | 15 | 282 | 1076 | Unknown | EF428256 | RGS3 | 4.566X10-03 |
|  | (CTT)4-x3-(AG)3 | 2520 | 2539 | 9 | 2219 | 2595 | MZ448274 | KR013745 | RMCS3 | 1.000X10-05 |
| FJ345399 | (CG)3-x1-(T)7 | 11 | 24 | 15 | 282 | 1076 | Unknown | EF428256 | RGS3 | 4.566X10-03 |
|  | (CTT)4-x3-(AG)3 | 2520 | 2539 | 9 | 2395 | 2583 | MZ448274 | KR013745 | RMCS3 | 1.000X10-05 |
| FJ345400 | (CG)3-x1-(T)7 | 11 | 24 | 15 | 467 | 1043 | Unknown | EF428256 | RGS3 | 4.566X10-03 |
|  | (CTT)4-x3-(AG)3 | 2520 | 2539 | 9 | 2395 | 2583 | MZ448274 | KR013745 | RMCS3 | 1.000X10-05 |
| FN645922 | (CG)3-x1-(T)9 | 11 | 26 | 8 | 1096 | 1458 | FJ159262 | FJ159267 | RMCS3 | 4.355X10-07 |
|  | (AT)3-x-1-(T)7 | 2462 | 2473 | 10 | 1629 | 1702 | FJ345398 | FJ159267 | RGMCS3 | 1.288X10-08 |
|  | (CTT)3-x4-(GA)3 | 2511 | 2529 |  |  |  |  |  |  |  |
| HE578897 | (CG)3-x2-(T)6 | 11 | 24 | 6 | 2516 | 2739 | MH628534 | FJ159265 | RGBMCS3 | 1.966X10-13 |
|  | (AT)3-x-1-(T)6 | 2470 | 2480 |  |  |  |  |  |  |  |
| JX181786 | (CG)3-x1-(T)9 | 11 | 26 | 8 | 1071 | 1458 | FJ159262 | FJ159267 | RMCS3 | 4.355X10-07 |
|  | (AT)3-x-1-(T)7 | 2462 | 2473 | 10 | 1629 | 1702 | FJ345398 | FJ159267 | RGMCS3 | 1.288X10-08 |
|  | (CTT)3-x4-(GA)3 | 2511 | 2529 |  |  |  |  |  |  |  |
| KJ462083 | (CG)3-x1-(T)8 | 11 | 25 | 10 | 1621 | 194 | FJ345398 | FJ159267 | RGMCS3 | 1.288X10-08 |
|  | (AT)3-x-1-(T)7 | 2461 | 2472 | 8 | 1095 | 1457 | FJ159262 | FJ159267 | RMCS3 | 4.355X10-07 |
|  | (CTT)3-x4-(GA)3 | 2510 | 2528 |  |  |  |  |  |  |  |
| KR013745 | (CG)3-x1-(T)9 | 11 | 26 | 13 | 328 | 458 | MH628534 | FJ345399 | RGBM3 | 1.471X10-03 |
|  |  |  |  | 11 | 2388 | 2689 | MZ448277 | Unknown | RMCS3 | 9.888X10-11 |
| KT948076 | (CG)3-x1-(T)7 | 11 | 24 | 13 | 329 | 459 | MH628534 | FJ345399 | RGBM3 | 1.471X10-03 |
|  |  |  |  | 12 | 1847 | 2239 | KR013745 | Unknown | RGMCS3 | 2.459X10-13 |
| MH538339 | (CG)3-x1-(T)8 | 11 | 25 | 2 | 1810 | 2738 | MZ448280 | Unknown | RGBMCS3 | 3.489X10-36 |
| MH628534 | (CG)3-x1-(T)8 | 11 | 25 | 11 | 2288 | 2680 | MZ448277 | Unknown | RMCS3 | 9.888X10-11 |
| MN005041 | (CG)3-x1-(T)8 | 11 | 25 | 8 | 1093 | 1457 | FJ159262 | FJ159267 | RMCS3 | 4.355X10-07 |
|  | (TC)3-x2-(CT)4 | 2612 | 2627 | 10 | 1620 | 1701 | FJ345398 | FJ159267 | RGMCS3 | 1.288X10-08 |
|  |  |  |  | 1 | 1836 | 2733 | MZ448280 | Unknown | RGBMCS3 | 1.000X10-39 |
| MZ448274 | (CG)3-x1-(T)7 | 11 | 24 | - | - | - | - | - | - | - |
|  | (AT)3-x-1-(T)7 | 2472 | 2483 |  |  |  |  |  |  |  |
| MZ448275 | (CG)3-x1-(T)7 | 11 | 24 | - | - | - | - | - | - | - |
|  | (AT)3-x-1-(T)7 | 2472 | 2483 |  |  |  |  |  |  |  |
| MZ448276 | (CG)3-x1-(T)7 | 11 | 24 | - | - | - | - | - | - | - |
|  | (AT)3-x-1-(T)7 | 2472 | 2483 |  |  |  |  |  |  |  |
| MZ448277 | (CG)3-x1-(T)7 | 11 | 24 | - | - | - | - | - | - | - |
|  | (AT)3-x-1-(T)7 | 2472 | 2483 |  |  |  |  |  |  |  |
| MZ448278 | (CG)3-x1-(T)7 | 11 | 24 | - | - | - | - | - | - | - |
|  | (AT)3-x-1-(T)7 | 2498 | 2509 |  |  |  |  |  |  |  |
| MZ448279 | (CG)3-x1-(T)7 | 11 | 24 | - | - | - | - | - | - | - |
|  | (AT)3-x-1-(T)7 | 2472 | 2483 |  |  |  |  |  |  |  |
| MZ448280 | (CG)3-x1-(T)7 | 11 | 24 | - | - | - | - | - | - | - |
|  | (AT)3-x-1-(T)7 | 2485 | 2496 |  |  |  |  |  |  |  |
| NC_009088 | (CG)3-x1-(T)7 | 11 | 24 | - | - | - | - | - | - | - |
|  | (AT)4-x-1-(T)8 | 2444 | 2458 |  |  |  |  |  |  |  |
| NC_010818 | (CG)3-x1-(T)7 | 11 | 24 | 3 | 1751 | 31 | Unknown | MZ448280 | RGBMCS3 | 2.372X10-35 |
|  | (AT)3-x-1-(T)7 | 2457 | 2468 |  |  |  |  |  |  |  |

**Note**. ^1^ R-RDP; G-GENCONV; B-BOOTSCAN; M-MAXCHI; C-CHIMERA; S-SISCAN; 3-3SEQ

^2^ The lowest p-value detected by one of the methods.

**Table S4.** Identification of the recombination breakpoints and cSSR motifs in betasatellite isolates

| **Accession number** | **cSSR analysis** | | | **Recombination analysis** | | | | | | |
| --- | --- | --- | --- | --- | --- | --- | --- | --- | --- | --- |
|  | **cSSR motif** | **cSSR position** | | **Event number** | **Recombination breakpoints** | | **Putative parental sequences** | | **RDP methods^1^** | ***p*-value^2^** |
|  |  | **Start** | **End** |  | **Start** | **End** | **Major** | **Minor** |  |  |
| DQ298137 | (GA)3-x3-(TTG)4-x-2-(TG)3 | 602 | 626 |  |  |  |  |  |  |  |
|  | (A)9-x1-(A)6 | 948 | 963 |  |  |  |  |  |  |  |
| EF614158 | (GA)3-x3-(TTG)4-x-2-(TG)3 | 602 | 626 | 4 | 740 | 831 | DQ298137 | NC_028130 | RGBMCS3 | 6.247 ✕ 10^-12^ |
|  |  |  |  | 3 | 1200 | 1311 | MZ448282 | KT390489 | RGBMCS3 | 2.935 ✕ 10^-14^ |
| EF614159 | (GA)3-x3-(TTG)4-x-2-(TG)3 | 602 | 626 | 4 | 740 | 831 | DQ298137 | NC_028130 | RGBMCS3 | 6.247 ✕ 10^-12^ |
|  | (A)9-x1-(A)6 | 952 | 967 | 3 | 1200 | 1286 | MZ448282 | KT390489 | RGBMCS3 | 2.935 ✕ 10^-14^ |
| EF614160 | (TA)6-x4-(TA)3 | 737 | 758 |  |  |  |  |  |  |  |
|  | (A)6-x1-(A)15 | 898 | 919 |  |  |  |  |  |  |  |
|  | (A)7-x1-(A)6 | 949 | 962 |  |  |  |  |  |  |  |
| EF614161 | (TA)3-x9-(A)8 | 765 | 787 | 11 | 23 | 265 | KT948077 | FJ159274 | RGMCS | 3.430 ✕ 10^-8^ |
|  | (A)6-x9-(A)8 | 902 | 924 |  |  |  |  |  |  |  |
|  | (A)9-x1-(A)6 | 958 | 973 |  |  |  |  |  |  |  |
| EF614162 | (GA)3-x3-(A)7 | 948 | 963 | 11 | 17 | 272 | KT948077 | FJ159274 | RGMCS | 3.430 ✕ 10^-8^ |
| EU557374 | (TGG)3-x3-(GAA)4 | 44 | 67 | 11 | 1329 | 212 | KT948077 | FJ159274 | RGMCS | 3.430 ✕ 10^-8^ |
|  | (A)6-x7-(A)16 | 905 | 933 |  |  |  |  |  |  |  |
| FJ159272 | (A)6-x7-(A)11 | 905 | 928 | 11 | 1329 | 212 | KT948077 | FJ159274 | RGMCS | 3.430 ✕ 10^-8^ |
| FJ159273 | (A)8-x7-(A)16 | 903 | 933 | 11 | 1329 | 212 | KT948077 | FJ159274 | RGMCS | 3.430 ✕ 10^-8^ |
| FJ159274 | (GA)3-x3-(TTG)4-x-2-(TG)3 | 602 | 626 | 3 | 1200 | 1286 | MZ448282 | KT390489 | RGBMCS3 | 2.935 ✕ 10^-14^ |
|  | (TA)3-x2-(TA)4-x0-(TG)3 | 740 | 761 |  |  |  |  |  |  |  |
|  | (A)9-x1-(A)6 | 952 | 967 |  |  |  |  |  |  |  |
| FJ159275 | (GA)3-x3-(TTG)4-x-2-(TG)3 | 602 | 626 | 10 | 889 | 1140 | FJ159274 | Unknown | RMC3 | 2.805 ✕ 10^-5^ |
|  | (A)9-x1-(A)6 | 952 | 967 | 3 | 1200 | 1286 | MZ448282 | KT390489 | RGBMCS3 | 2.935 ✕ 10^-14^ |
| MZ448281 | (GA)3-x3-(TTG)4-x-2-(TG)3 | 601 | 625 |  |  |  |  |  |  |  |
|  | (A)9-x1-(A)6 | 947 | 962 |  |  |  |  |  |  |  |
| MZ448282 | (GA)3-x3-(TTG)4-x-2-(TG)3 | 601 | 625 | 15 | 1173 | 1286 | MZ448285 | Unknown | RGMCS3 | 6.311 ✕ 10^-8^ |
|  | (A)9-x1-(A)6 | 947 | 962 |  |  |  |  |  |  |  |
| MZ448283 | (GA)3-x2-(T)6-x5-(TG)3 | 616 | 640 |  |  |  |  |  |  |  |
|  | (A)8-x1-(A)6 | 974 | 988 |  |  |  |  |  |  |  |
| MZ448284 | (GA)3-x3-(TTG)4-x-2-(TG)3 | 601 | 625 |  |  |  |  |  |  |  |
|  | (A)9-x1-(A)6 | 946 | 961 |  |  |  |  |  |  |  |
| MZ448285 | (GA)3-x3-(TTG)4-x-2-(TG)3 | 601 | 625 |  |  |  |  |  |  |  |
|  | (TA)3-x4-(TA)3 | 740 | 755 |  |  |  |  |  |  |  |
|  | (A)8-x1-(A)6 | 947 | 961 |  |  |  |  |  |  |  |
| MZ448286 | (GA)3-x3-(TTG)4-x-2-(TG)3 | 602 | 626 |  |  |  |  |  |  |  |
| NC_009903 | (GA)3-x3-(TTG)4-x-2-(TG)3 | 602 | 626 |  |  |  |  |  |  |  |
|  | (A)9-x1-(A)6 | 948 | 963 |  |  |  |  |  |  |  |
| NC_010569 | (TGG)3-x3-(GAA)4 | 44 | 67 | 11 | 1329 | 212 | KT948077 | FJ159274 | RGMCS | 3.430 ✕ 10^-8^ |
|  | (A)6-x7-(A)16 | 905 | 933 |  |  |  |  |  |  |  |
| KT390489 | (TGG)3-x3-(GAA)4 | 45 | 68 | 2 | 1328 | 104 | Unknown | NC_028130 | RGBMCS3 | 2.680 ✕ 10^-9^ |
|  | (A)9-x1-(A)8 | 788 | 805 | 8 | 995 | 1198 | Unknown | EU880233 | RGBMCS | 1.901 ✕ 10^-14^ |
|  | (A)6-x8-(TC)4 | 1183 | 1204 | 5 | 1201 | 1320 | Unknown | KT390482 | RGBMCS3 | 1.051 ✕ 10^-13^ |
| KT390486 | (TGG)3-x3-(GAA)4 | 44 | 67 | 11 | 17 | 128 | KT948077 | FJ159274 | RGMCS | 3.430 ✕ 10^-8^ |
|  | (A)6-x6-(A)6 | 958 | 975 | 6 | 128 | 303 | EF614161 | Unknown | RBMCS3 | 1.016 ✕ 10^-10^ |
| KT390482 | (TGG)3-x3-(GAA)4 | 44 | 67 | 2 | 1207 | 100 | Unknown | NC_028130 | RGBMCS3 | 2.680 ✕ 10^-9^ |
|  | (TA)3-x9-(A)8-x4-(A)9-x1-(A)8-x9-(AAAGA)4 | 773 | 847 | 1 | 104 | 796 | Unknown | EF614161 | RGBMCS3 | 5.941 ✕ 10^-28^ |
|  | (A)7-x2-(A)12 | 922 | 942 |  |  |  |  |  |  |  |
|  | (A)8-x1-(A)10 | 970 | 988 |  |  |  |  |  |  |  |
| KT390479 | (TGG)3-x3-(GAA)4 | 44 | 67 | 11 | 1326 | 211 | KT948077 | FJ159274 | RGMCS | 3.430 ✕ 10^-8^ |
|  | (A)6-x9-(A)8 | 902 | 924 |  |  |  |  |  |  |  |
| NC_028130 | (TGG)3-x3-(GAA)4 | 44 | 67 | 11 | 17 | 128 | KT948077 | FJ159274 | RGMCS | 3.430 ✕ 10^-8^ |
|  | (A)6-x6-(A)6 | 958 | 975 | 6 | 128 | 303 | EF614161 | Unknown | RBMCS3 | 1.016 ✕ 10^-10^ |
| EF620566 | (GA)3-x5-(TGT)5 | 601 | 625 | 14 | 1119 | 66 | Unknown | MG587941 | RBMC3 | 2.006 ✕ 10^-4^ |
|  | (A)9-x1-(A)6 | 934 | 949 |  |  |  |  |  |  |  |
| EU431115 | (TA)3-x6-(AT)3-x4-(AT)3 | 587 | 614 |  |  |  |  |  |  |  |
|  | (CA)3-x5-(GAA)4-x-2-(A)7-x-12-(GAAA)6 | 918 | 957 |  |  |  |  |  |  |  |
| EU825206 | (GA)3-x5-(TGT)5 | 601 | 625 | 14 | 1119 | 1342 | Unknown | MG587941 | RBMC3 | 2.006 ✕ 10^-4^ |
|  | (A)7-x1-(A)6-x1-(A)7 | 936 | 957 |  |  |  |  |  |  |  |
| EU825205 | (GA)3-x5-(TGT)5 | 601 | 625 | 14 | 1118 | 131 | Unknown | MG587941 | RBMC3 | 2.006 ✕ 10^-4^ |
|  | (A)9-x1-(A)6 | 934 | 949 |  |  |  |  |  |  |  |
| EU880233 | (TA)3-x6-(AT)3-x4-(AT)3 | 587 | 614 |  |  |  |  |  |  |  |
|  | (CA)3-x6-(A)8-x1-(A)7-x-12-(GAAA)6 | 918 | 957 |  |  |  |  |  |  |  |
| EU880232 | (TA)3-x6-(AT)3-x4-(AT)3 | 587 | 614 |  |  |  |  |  |  |  |
|  | (CA)3-x5-(GAA)4-x-2-(A)7-x-12-(GAAA)6 | 918 | 957 |  |  |  |  |  |  |  |
| EF620564 | (GA)3-x3-(TTG)4-x-2-(TG)3 | 601 | 625 |  |  |  |  |  |  |  |
|  | (A)8-x1-(A)6 | 947 | 961 |  |  |  |  |  |  |  |
| EU880231 | (GA)3-x3-(TTG)4-x-2-(TG)3 | 601 | 625 |  |  |  |  |  |  |  |
|  | (A)8-x1-(A)6 | 947 | 961 |  |  |  |  |  |  |  |
| EF620565 | (GA)3-x3-(TTG)4-x-2-(TG)3 | 601 | 625 |  |  |  |  |  |  |  |
|  | (A)14-x8-(CGT)3 | 904 | 934 |  |  |  |  |  |  |  |
|  | (A)7-x1-(A)6 | 949 | 962 |  |  |  |  |  |  |  |
| MK358823 | (GA)3-x3-(TTG)4-x-2-(TG)3 | 601 | 625 | 7 | 1165 | 1334 | MZ448286 | EU557374 | RGM3 | 6.088 ✕ 10-8 |
| NC_027133 | (TTG)4-x-2-(TG)3 | 612 | 627 |  |  |  |  |  |  |  |
|  | (A)8-x1-(A)6 | 937 | 951 |  |  |  |  |  |  |  |
| FR772083 | (TTG)4-x-2-(TG)3 | 612 | 627 |  |  |  |  |  |  |  |
|  | (A)8-x1-(A)6 | 937 | 951 |  |  |  |  |  |  |  |
| FN678779 | (TTG)4-x-2-(TG)3 | 612 | 627 |  |  |  |  |  |  |  |
|  | (A)8-x1-(A)6 | 937 | 951 |  |  |  |  |  |  |  |
| LK028572 | (TTG)4-x-2-(TG)3 | 611 | 626 |  |  |  |  |  |  |  |
|  | (A)17-x4-(A)14 | 870 | 904 |  |  |  |  |  |  |  |
| KT948077 | (A)9-x6-(A)15 | 877 | 906 |  |  |  |  |  |  |  |
|  | (A)9-x1-(A)6 | 935 | 950 |  |  |  |  |  |  |  |
|  | (GTAA)3-x9-(A)6 | 1022 | 1048 |  |  |  |  |  |  |  |
| LT716981 | (TGT)3-x9-(GT)3 | 145 | 169 | 9 | 1107 | 1222 | Unknown | EF614161 | RGMCS3 | 3.132 ✕ 10^-7^ |
| MG587941 | (T)6-x3-(A)22 | 890 | 920 |  |  |  |  |  |  |  |
| MG587933 | - |  |  | 20 | 776 | 896 | MG587941 | Unknown | RMC3 | 2.249 ✕ 10^-4^ |
| MF683831 | (TTG)4-x-2-(TG)3 | 612 | 627 | 13 | 499 | 939 | Unknown | FR772083 | RGBMCS3 | 1.706 ✕ 10^-8^ |
|  | (A)9-x1-(A)7 | 934 | 950 |  |  |  |  |  |  |  |
| MF409173 | (TTG)4-x-2-(TG)3 | 614 | 629 |  |  |  |  |  |  |  |

**Note**. ^1^ R-RDP; G-GENCONV; B-BOOTSCAN; M-MAXCHI; C-CHIMERA; S-SISCAN; 3-3SEQ

^2^ The lowest p-value detected by one of the methods.
